# BOWIe: Wireless Headstage for Optical Stimulation in Closed-Loop Optogenetics Experiments in Freely-moving Rodents

**DOI:** 10.64898/2026.10.13.758187

**Authors:** Patrícia Silva, Margarida Falcão, Sara L. Reis, Francisco Neves, Mafalda Abrantes, Tiago Pereira, Patricia Monteiro, Luis Jacinto

**Author notes:** Department of Biomedicine – Experimental Biology Unit, Faculty of Medicine of the University of Porto (FMUP), 4200-319, Porto, Portugal, *Email addresses* (L. Jacinto).

## Abstract

Optogenetics is an invaluable technique in neuroscience to study brain circuits, but conventional optogenetic systems use tethered approaches to deliver light to the brain which can hinder natural behavior. Optoelectronic advances led to the development of wireless optical stimulators for optogenetics experiments, but most reported devices have low widespread use due to complex builds and reduced versatility, which has also limited their integration in closed-loop systems. Here, we developed BOWIe, a battery-powered, low-weight, wireless headstage for photostimulation in optogenetics experiments with rodents, especially designed for simple integration in multiple behavioral setups, including those requiring closed-loop operation. BOWIe is based on a Bluetooth-Low Energy system-on-a-chip that allows simultaneous control and communication capabilities with a reduced number of components, and can be paired with an Arduino-based transmitter for simple integration in any behavioral paradigm using analogue or digital triggers. Two modes of optical stimulation are possible, either with ferrule-coupled optical fibers or µLED neural probes, which permit adapting the headstage according to material availability or experimental demands. A user-friendly app was also developed to simplify headstage pairing, definition of stimulation parameters, and protocol design. BOWIe was validated in two behavioral paradigms, including in a closed-loop experiment using computer vision, Bonsai RX, and Deeplabcut Live to track behavioral parameters online and remotely trigger photostimulation upon detected events. BOWIe’s modular design and flexible integration with behavioral setups make it a practical tool for in vivo experiments with optogenetics.

## 1. Introduction

Optogenetics is an ingenious technique combining genetic manipulation and optics to allow precise temporal and spatial control of neuronal activity through the optical activation of light-gated ionic channels expressed in neuronal membranes [1], [2]. Optogenetics has been fundamental to the investigation of brain circuitry and has contributed to the advancement of our knowledge on how brain disorders affect brain circuits and activity [2]–[4]. For optical stimulation in vivo, where causal relations between the brain area or cell type activated and behavior can be studied, light needs to be delivered to the brain [5]. Conventional approaches to deliver optical stimulation in behavioral experiments with rodents typically involves using a laser or a high-brightness LED patched to an optical fiber implanted on the animal’s brain region of interest [6], [7]. However, tethered experiments may hinder natural behavior and induce distress in animals due to tether entanglements leading to head tension. Particular experimental paradigms, such as social interaction studies [8], [9] and maze exploring tasks [10], also benefit from wire-free instrumentation.

To overcome the limitations of tethered optical delivery to the brain, wireless devices for optogenetic stimulation in rodents emerged as a potential solution, and different wireless headstages have been proposed in recent years [11]. These devices can be controlled remotely through wireless communication, including radio-frequency (RF) [12]–[18], infrared radiation (IR) [19], electromagnetism [20], or Bluetooth [21], [22], and can be powered by batteries [13], [16], [17], [19]–[21], [23], [24] or inductive RF coupling [9], [14], [15], [18], [25]–[27]. In terms of optical delivery, the proposed headstages focused on one type of light delivery mode to the brain, either through ferrule-coupled optical fibers [19], [20], [28] or, most commonly implanted, µLEDs [9], [14], [16], [22]. The use of modular headstages that would allow using both optical fibers and implantable µLEDs probes has not yet been reported.

Despite the progressive advance and refinement of these wireless headstages, most devices are still limited to short working ranges due to the need for close proximity and alignment for inductive powering and low optical power output [11]. Additionally, many of these devices cannot be easily reproduced by other labs due to complex or lacking build instructions, the absence of user-friendly apps for device programming and control, or because they have been designed for specific experimental setups, limiting their widespread use. This includes their limited integration in closed-loop systems, in which behavioral or physiological biomarkers can be tracked in real-time to trigger brain stimulation according to detected events. Although these systems are fundamental to advancing causal investigation of brain activity modulation, reported systems have focused on tethered approaches only [29], [30].

To address current limitations and permit the simple integration of wireless control of optical stimulation in closed-loop behavioral paradigms, we developed BOWIe, a Brain Optogenetics Wireless Interface headstage, that takes advantage of a BLE system-on-a-chip (SoC) to reduce size and weight and simplify operation. BOWIe can be assembled with commercially available off-the-shelf components and communicates with a user app via an Arduino-compatible emitter that can be integrated in closed-loop paradigms with analogue or digital stimulation triggers. BOWIe also has a modular design that permits the use of either ferrule-coupled optical fibers or µLED neural probes for flexible choice of optical delivery mode, according to experimental demands or user preference. Both uses were validated in behavioral paradigms, including in a closed-loop experiment in which optical stimulation was wireless triggered via BOWIe from specific behavioral events detected with DeepLabCut running on Bonsai-RX. Overall, these findings establish BOWIe as a versatile and effective platform for wireless optogenetic stimulation in rodent experiments, facilitating the integration of wireless stimulation with closed-loop paradigms.

## 2. Results and Discussion

### 2.1. System description, implementation, and operation

To permit wirelessly controlled optical stimulation in optogenetics experiments with rodents, we developed BOWIe, a compact Brain Optogenetics Wireless Interface headstage especially designed for closed-loop behavioral paradigms (Figure 1). BOWIe was built around a Bluetooth-Low-Energy (BLE) System-on-a-chip (SoC) (CYBLE-014008, Infineon Technologies) to reduce circuitry complexity and minimize system size. The wire-free headstage is composed of four main parts (Figure 1A): a printed-circuit board (PCB) to hold and connect the essential electronic components, a compact 30 mAh lithium polymer (LiPo) battery for power supply, the BLE SoC for control and communication, and a modular interface for optical light emission with fiber-coupled ferrules or µLED probes (Figure 1B). Besides the BLE SoC and the battery, the following components are mounted on the PCB: an indicator LED to provide optical feedback on the system’s state, a 3.5 V voltage regulator to ensure stable power from the battery to the optical emitter, a 330 Ω surface-mount ferrite bead to filter noise from the battery to the BLE SoC, and a dip-switch for turning the circuit on and off (Figure 1A). The SoC also uses an integrated ADC to read battery voltage and controls the indicator red LED on the top part of the headstage.

**Figure 1.**
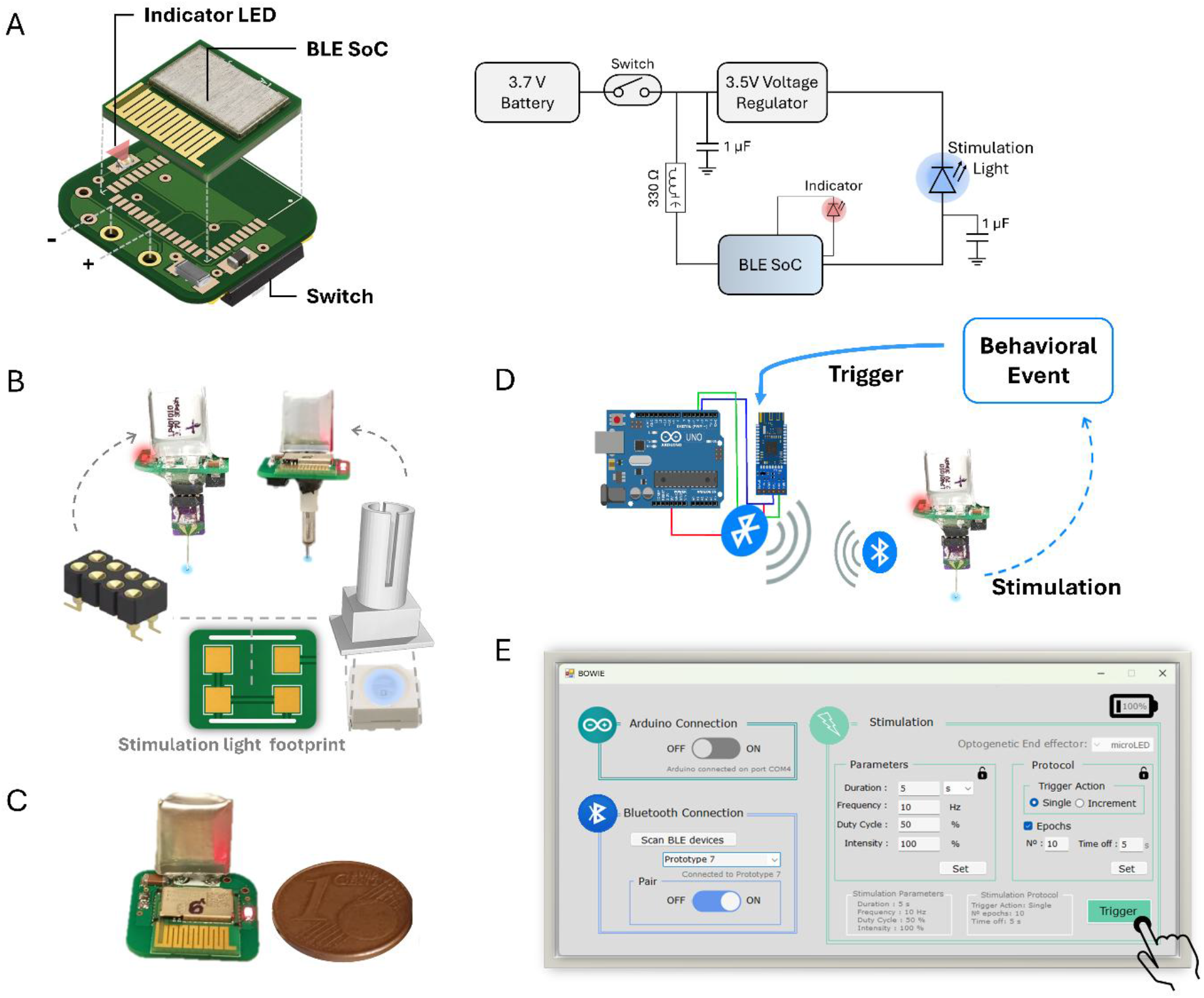
Overview of BOWIe headstage. A – Printed circuit board (PCB) mounting configuration (left) and schematic of the stimulator circuit (right), using a BLE SoC, a 3.7V battery, a switch, a 3.5V voltage regulator, an indicator LED, a ferrite bead, and capacitors; B – Schematic of the two interchangeable coupling interfaces for either a µLED neural probe or an optical fiber, with the different mounting scheme at footprint on the PCB bottom view, including a Millmax connector for coupling with a µLED neural probe or a surface mount LED and a 3D-printed resin piece for coupling with an optical fiber; C – Assembled version of BOWIe showcasing its small size; D – Conceptual diagram of BOWIe’s closed-loop operation, illustrating the link between behavioral event detection, wireless Bluetooth communication and triggering, and photostimulation; E – Custom designed app for device pairing, stimulation parameter control, protocol design, manual triggering and battery assessment.

The headstage has a modular structure to allow the wireless control of photostimulation through both implantable ferrule-coupled optical fibers or µLED neural probes (Figure 1B) according to experimental needs or material availability. Thus, to use an optical fiber for photostimulation, the headstage is assembled with a surface mount LED connected to the bottom part of the PCB (Figure1B). A custom 3D printed resin piece then serves as a coupler between the LED and the optical fiber ferrule to ensure light collimation alignment. To use the headstage with our previously designed implantable µLED neural probes [31], a Millmax pin connector must be soldered to the bottom part of the headstage, instead of the surface mount LED. A fully assembled headstage is shown on Figure 1C. BOWIe was designed to pair with an Arduino-compatible Bluetooth module (HM-10 BLE, CN0022, SunFounder) (Figure 1D) to facilitate integrating different analogue or digital trigger commands in different types of experimental designs. The Arduino code handles the communication interface between the headstage and the commands from a custom-designed app (Figure 1E).

A standalone app with a user-friendly interface was designed to facilitate device connection and programming stimulation parameters to users without programming skills (Figure 1E). The app is divided into 4 main components: computer communication and Bluetooth pairing, battery assessment, stimulation parameters definition, and type of triggering mode. Once launched, the app automatically scans the PC’s USB ports for connected Arduinos running the program for the HM-10 BLE transmitter. The user can then press a button for automatic scanning of available Bluetooth devices allowing the selection of the headstage and its pairing with the HM-10 BLE transmitter. Once connected, the user can assess the battery status via the app, which provides real-time feedback based on voltage measurements taken from an ADC in the BLE chip connected to the battery. Depending on the photostimulation method used with the headstage, the user must select in the app the desired optical delivery method, so that the app can automatically regulate the maximum output voltage to either 3.5 V or 3.2V for the LED-optical fiber or μLED neural probe, respectively The user can finally define stimulation duration (in seconds or minutes), frequency, duty cycle, and intensity (in percentage of maximum intensity). The app also supports protocol design, such as setting the number of photostimulation epochs and the time off between them. After configuration of the stimulation parameters, the stimulation protocol can be triggered at the headstage by transistor-to-transistor-logic (TTL) inputs to the Arduino with the Bluetooth transmitter (default mode) or with the pressing of a trigger button on the app. The use of TTL inputs to the Arduino as external triggers allows simple integration of this system into multiple commercial and custom-designed behavioral setups, which typically already use TTL signals, extending their operation to closed-loop experiments.

Table 1 compares BOWIe with previously available wireless headstages for optogenetics experiments in rodents. Distinctive features of BOWIe include the option of using two different modes for optical stimulation delivery, higher irradiances when the µLED neural probe is used, a higher working distance, and the presence of a user interface app. Of note, although Bluetooth chips have been used in previous wireless headstages for optical stimulation for rodents [21], [22], albeit less frequently than other options with RF communication, the integration of a BLE SoC had not been reported. The BLE SoC controls not only communication but also makes use of an integrated DAC to provide the necessary stable voltage to drive the light delivery system (surface mount LED or µLED in probe).

**Table 1.**
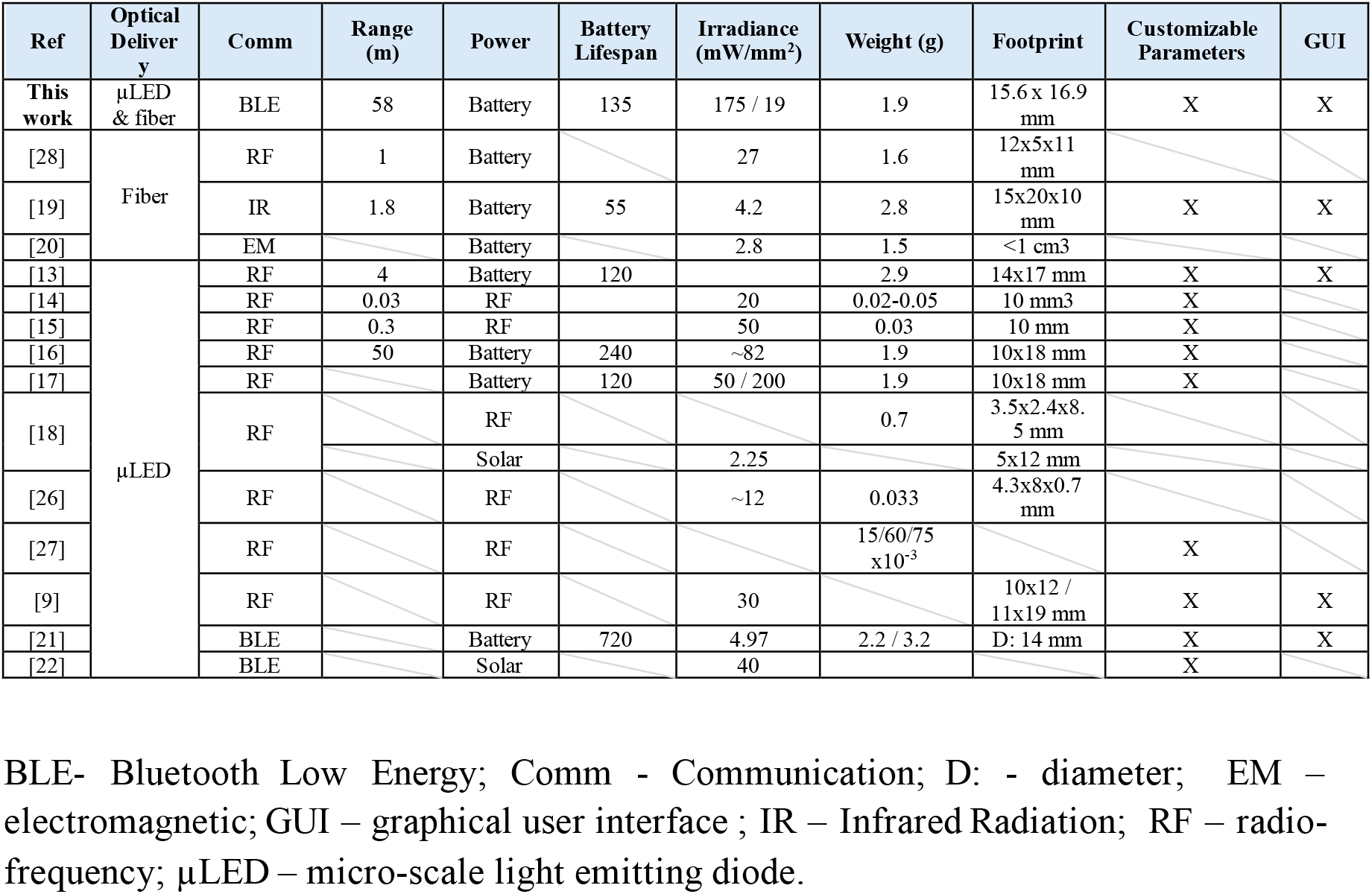
Wireless optical stimulators for freely moving mice optogenetics experiments.

### 2.2. System characterization

#### 2.2.1. Device assembly

BOWIe can be assembled with off-the shelf electronic components soldered to a PCB. PCB fabrication files and the list of components, as well as assembly and programming instructions, are publicly hosted on a GitHub repository (https://github.com/neuroephys-lab/BOWIe).

#### 2.2.2. Weight and size

The device PCB’s footprint measures 15.60 x 16.90 x 0.80 mm, with the 10 x 10 x 4 mm battery adding an additional 10 mm in height. The total weight of the headstage with battery, but without the optical delivery system, is 1.66 g, which increases to 1.90 g and 1.99 g for the optical fiber and µLED neural probe versions, respectively. This weight is below the widely accepted limit of 3 g (or 10% of adult weight) for headstages for mice. When compared to other wirelessly controlled headstages for optical stimulation (Table 1), BOWIE presents a footprint and a weight that is comparable to the lightest devices using batteries.

#### 2.2.3. Optical delivery mode

The amount of optical power delivered to the brain can have a significant impact on optogenetic experimental outcomes, as insufficient light will not trigger neuronal activation or inhibition [32] and excessive light converted to heat can cause neuronal damage or unwanted modulation of activity [33]. When using the ferrule-coupled optical fiber approach with BOWIe there is a significant loss of optical power between the surface mount LED on the headstage and the optical fiber due to the uncollimated light emission and refraction at the interface. When driving the LED at maximum intensity, the 40 mW optical power output from the LED is reduced to 0.6 mW at the tip of a 200 µm fiber with a numerical aperture (NA) of 0.22. Nevertheless, the irradiance at the tip was calculated to be approximately 19 mW/mm^2^, which is above the limit reported to trigger neuronal activity in neurons with channelrhodopsin [34]. When comparing the obtained irrandiance with a previously reported fiber-coupled wireless stimulator [19], BOIWe presents irradiance values approximately 4.5 times higher using a 200 µm fiber instead of 500 µm fiber.

For the µLED neural probe option, BOWIe can output up to 2.5 mW with an irradiance of 175 mW/mm^2^ [26] when driven at maximum intensity. But due to heating of the µLED surface, stimulation protocols at 10 or 20 Hz have to be limited to 20% duty cycles for any output amplitude, or limited to approximately 80% of the amplitude for duty cycles between 20 and 80% [31]. The latter still corresponds to a maximum optical power of approximately 0.80 mW with an irradiance of 40 mW/mm^2^. Altough both optical delivery options will work to modulate neuronal activity with BOWIe, the implantable version makes more efficient use of the device supply power by increasing the amount of battery output power that is converted into optical power.

#### 2.2.4. Battery runtime and operating range

A challenge in designing a battery-powered stimulator lies in minimizing the current consumption, as a higher energy demand decreases the battery lifespan. This issue was addressed in BOWIe by using the sleep-mode functionality of the BLE communication protocol, so that the current draw was minimal while on stand-by, i.e. waiting for a trigger to deliver optical stimulation. BOWIe runs on a compact 30 mAh rechargeable LiPo battery, which was chosen because of its small footprint and weight while still showing a relatively high capacity for its size. The battery runtime during BOWIe operation was assessed for stand-by mode (with the device paired with the Bluetooth emitter but not delivering light output). In this state most of the battery power is consumed to keep the device in sleep mode and powering the small surface mount indicator LED on the PCB. This LED operates in three distinct operational states: when the headstage is turned on the LED blinks rapidly for 1 s and then turns off; when the BLE module is unpaired, the LED remains off; and when the module is paired, the LED remains on. This allows users to visually track if pairing is maintained throughout experimental conditions. When paired in stand-by mode, BOWIe achieved a maximum runtime of 135 minutes.

To evaluate battery duration during experiments with optical stimulation, runtime was assessed during continuous 10 Hz stimulation at maximum intensity and with intermittent (10 seconds ON, 10 seconds OFF) 10 Hz stimulation with the optical fiber and µLED probes. (Figure 2A). During the continuous 10 Hz stimulation with the optical fiber, the maximum battery runtime was 37 minutes, increasing to 60 minutes when intermittent stimulations was used. The battery maximum operating runtime with the µLED driven continuously at 10 Hz was 69 minutes, increasing to 116 minutes with intermittent stimulation. The later was expected as the µLED draws considerably less the current than the LED light source for the optical fiber stimulation. For practical experiments, where stimulation is not continuously applied, battery runtime is expected to increase further for both fiber and µLED optical stimulation because of the integration of the sleep-mode functionality.

**Figure 2.**
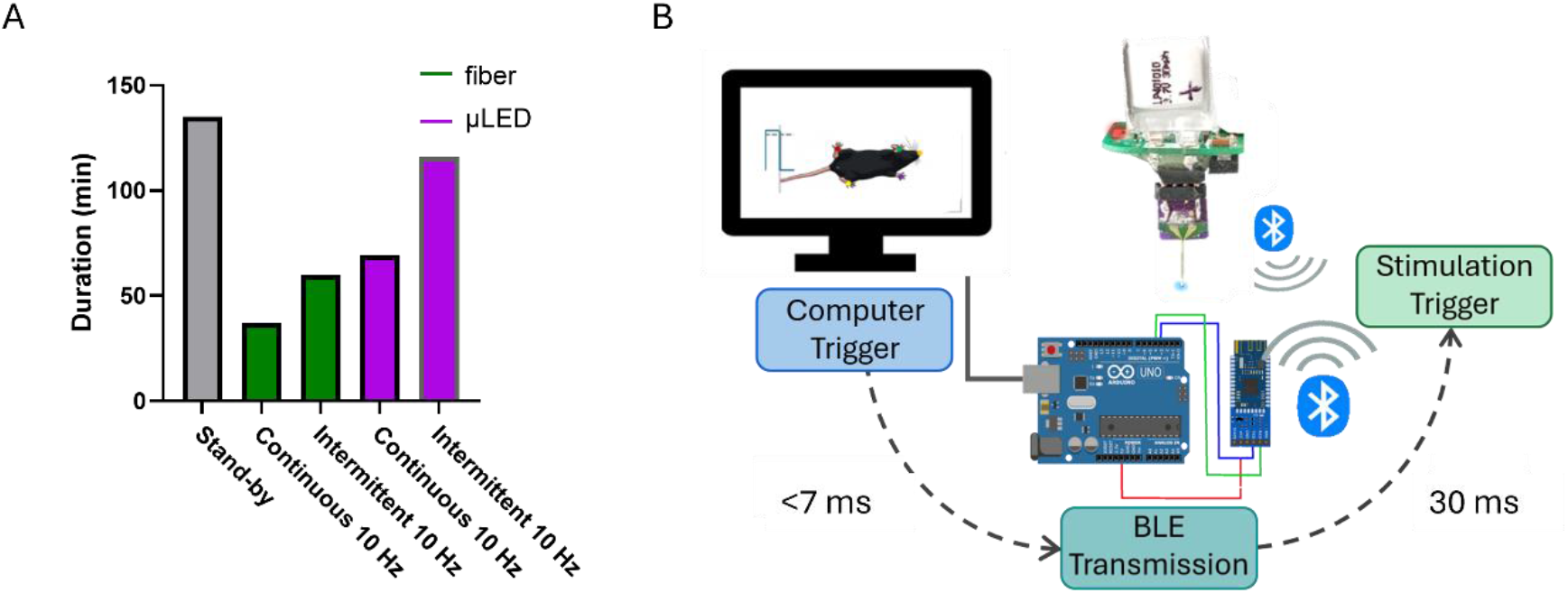
Characterization of BOWIe’s battery runtime and system delays. A) Battery runtime for standy mode (grey) and in different stimulation protocols including continuous 10 Hz and intermittent (10 s ON / 10 a OFF) stimulation for BOWIE with optical fibers (green) and µLED neural probe (violet); B) Schematic of BOWIE’s communication with the Arduino transmitter and a computer running a behavioral task, including system delays between a computer trigger and the Arduino and between BLE transmission and the stimulation turning ON.

The distance operating range of wireless headstages is important because it defines the maximum distance at which the emitter or computer must be from the headstage to operate which can impose limits on arena size and proximity of instrumentation or researchers. BOWIe’s maximum transmission range is approximately 58 meters without walls, which is the highest reported for wireless opto-stimulators (Table 1). This long range enables the positioning of the communication station far away from the animal’s experiment, reducing possible electromagnetic, noise and light interferences. Additionally, it may also enable more naturalistic paradigms with large or multi-room environments.

#### 2.2.5. System latency

The key advantage of closed-loop systems lies in their ability to act on variables in real time, thus the system’s response time is a crucial parameter to ensure a causal link between the detected event and the applied stimulation. To optimize BOWIe’s online performance, any pre-processing of parameters necessary to deliver an optical stimulus, or stimulation protocol, occurs at the app level and not on the BLE chip. This ensures that when receiving a trigger, the BLE chip on the headstage can immediately turn on the stimulation with the desired stimulation parameters or protocol. Using visual cues along the system’s path, the latency from the host computer trigger to the start of BLE transmission toward the receiver chip was measured to be below 7 ms. The delay between the stimulation request and the actual onset of stimulation was approximately 30 ms (Figure 2B). When compared with previous reports of wired optogenetics closed-loop systems, our latency value is within the lowest reported [29], [30].

### 2.3. Behavioral validation

#### 2.3.1. Photostimulation control in BOWIe with optical fiber

To validate the operation of our wirelessly controlled headstage to induce measurable behavioral changes during photostimulation with optical fiber ferrule implants, transgenic mice carrying channelrhodospin in the cortex (Emx1:Ai27D) [35], [36] were implanted with optical fibers in the secondary motor cortex (M2) of the left hemisphere (Figure 3A). BOWIe headstage was connected to the fiber ferrules and mice were allowed to freely explore an open arena. Following a 5 min habituation period, stimulation at 20 Hz with a 20% duty cycle was turned on and off in 1 minute epochs, for a total of 5 ON and 5 OFF epochs (Figure 3B). During photostimulation, an increase of contralateral rotations was observed when compared with ipsilateral rotations as well as with contralateral and ipsilateral rotations during periods with no stimulation (Figure 3C). Concomitantly, an increase in distance travelled during photostimulation periods was observed, most probably linked to the increase in the number of contralateral rotations (Figure 3D). The increase of both locomotor activity and contralateral rotations during M2 stimulations have been previously described [37], [38]. This confirms that photostimulation with optical fibers controlled by BOWIe can reliably induce expected behavioral changes, demonstrating the effectiveness of our headstage for tether-free optogenetics experiments.

**Figure 3.**
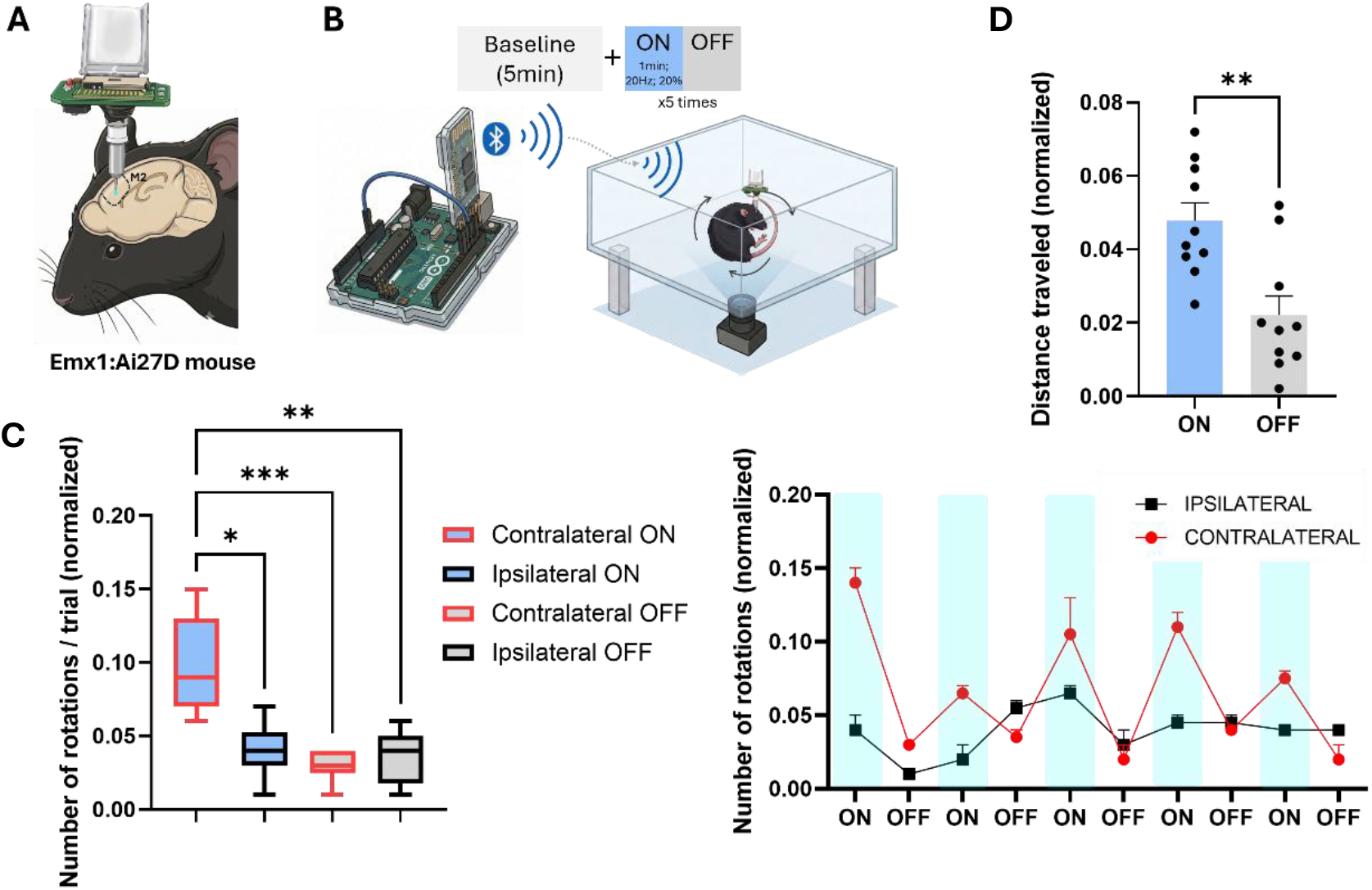
Wireless optogenetic stimulation of M2 brain region with optical fibers. **(A)** Schematic of the optic fiber implantation targeting the secondary motor cortex (M2) of an Emx1:Ai27D mouse. **(B)** Experimental design illustrating the protocol programmed in BOWIe, with a 5 minute baseline period followed by 5 stimulation period of 1 minute each with 1 minute trial intervals **(C)** Box-and-whisker plots comparing the normalized number of contralateral and ipsilateral rotations during stimulation ON and OFF states (left) and temporal profile of rotational responses (ipsilateral vs. contralateral) across consecutive stimulation blocks (light blue shaded regions indicate ON periods). **(D)** Total distanced travelled during the ON and OFF periods of optostimulation (470nm, 20Hz, 20% duty cyle).

#### 2.3.2. Closed-loop photostimulation with BOWIe with µLED neural probe

To evaluate closed-loop operation in vivo with our headstage with µLED neural probes, the ventrolateral periaqueductal gray (vlPAG), a midbrain structure known to mediate defensive freezing behaviour, was targeted (Figure 4A). A closed-loop system with a custom Bonsai RX [39] script using DeepLabCut (DLC) Live [40] to continuously track the animal’s body parts was employed to trigger µLED photostimulation during active locomotion to induce freezing (Figure 4B). During an initial 5 minute baseline period, without optical stimulation, the mouse exhibited a standard exploratory pattern (Fig. 4C, D). After the baseline period, 10 Hz photostimulation of the vlPAG was activated for 10 seconds, with at least 30 seconds intervals, whenever translational movement was detected by an algorithm running in Bonsai RX that provided online behavioral classification based on the animals’ body partes estimated by DLC-Live. Additional algorithms running online in Bonsai RX automatically detected and counted freezing and grooming events. A significant decrease of movement during stimulation with a progressive reduction in the mouse locomotor activity was observed (Figure 4C,D). This was accompanied by a reduction of the the time spent on grooming behavior (Figure 4E), an increase of freezing behavior (Figure 4F), and a reduction of distance travelled (Figure 4G when comparing the baseline period and the photostimulation periods. Quantitative analysis of the photostimulation epochs revealed a significant decrease in mean speed during the 10-second stimulation window compared to the 5-second pre-stimulation period (Fig. 4H). These findings are consistent with previous studies showing that stimulation of the vlPAG induce freezing behavior in mice [41], [42], particularly when using low-intensity stimulations [43]. Collectively, this behavioral response provides functional validation that BOWIe is capable of robustly modulating neuronal circuits in vivo in a closed-loop paradigm.

**Figure 4.**
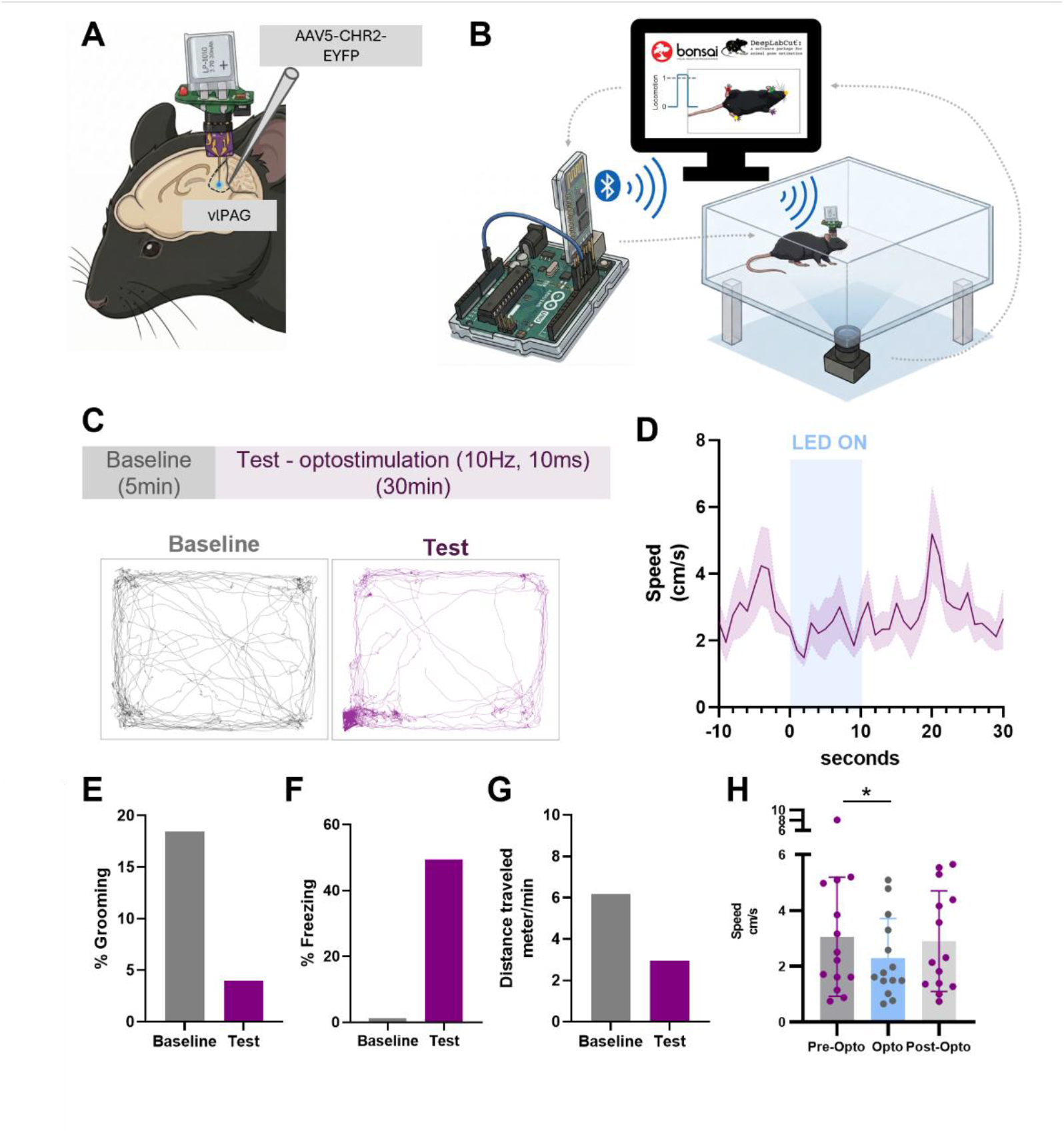
In vivo optogenetic modulation using µLED probes. (A) Schematic of the µLED probe implantation targeting the vlPAG. (B) Closed-loop optogenetic stimulation protocol. (C) Representative spatial occupancy plots of the mouse during the baseline (left; gray) and test (right; purple) session. (D) Peristimulus time histogram centered on µLED activation onset (t =0, 470nm, 10Hz, 10s duration)). Percentage of time spent grooming (E) and freezing (F) during baseline and test periods. (G) Total distance traveled during baseline vs test. (H) Quantitative analysis of mean speed across pre-stimulation (pre-opto), stimulation (opto), and post-stimulation (post-opto) epochs. *p < 0.05, paired *t*-test.

## 3. Conclusions

Here we presented BOWIe, a headstage for wireless control of optical stimulation in rodent experiments. BOWIe was specially designed with versatility in mind, so that it could be used according to experimental demands and a wide range of experimental setups. For this, both optical fibers and µLED neural probes can be used with BOWIe for optical delivery inside the brain, and any experimental setup using TTL triggers can be combined with BOWIe for closed-loop operation. Additionally, a user app was designed to facilitate pairing the device with the transmitter and allow the remote control of various stimulation parameters (duration, frequency, duty cycle and intensity) and protocol design.

BOWIe was validated in two behavioral setups, using either optical fibers or µLED neural probes, including in one example with closed-loop operation in which behavioral parameters were tracked in real-time to remotely trigger optical stimulation upon the detection of events of interest. The integration of BOWIe with other toolboxes frequently used in neuroscience experiments such as Bonsai RX and Deeplabcut was also demonstrated in this experiment.

One limitation of BOWIe is its dependence on battery power which reduces experiments to the battery runtime, and is constrained by type of stimulation delivery mode and protocol used. On the other hand, battery power allows the long-rage operation of BOWIe by breaking free of spatial constraints needed for inductive powering. Another limitation is that only one headstage can be controlled at a time, which limits BOWIE’s use on multi-animal paradigms or running parallel experiments without the need to replicate the full system. Nevertheless, as BOWIe is validated on more behavioral paradigms and different labs, its proven flexibility will contribute to the development of robust causal investigation of brain activity modulation in neuroscience optogenetics experiments.

## 4. Materials and Methods

### 4.1. Headstage design and assembly

A custom-designed PCB (15.61×16.89×0.8mm, Autodesk Fusion) with ENIG finishing was fabricated to assemble the headstage circuit. This circuit included a BLE system-on-a-chip (SoC) chip (CYBLE-014008, Infineon Technologies), a 330 Ω ferrite bead (BLM21PG331SN1D, Murata Electronics), two 1 µF capacitors (CC1206MRY5V9BB105, YAGEO), a toggle switch (113-DMR01V, Diptronics), a 3.5 V voltage regulator (NCP161ASN350T1G, onsemi), a 3.7 V rechargeable lithium-polymer battery (lp401010, LiPoly Batteries), and an indicative red LED (755-CSL1901VW1, ROHM Semiconductor). The modular bottom part of the PCB has a footprint for soldering either a blue surface-mount LED (150141BS73140, Wurth Elektronik) for fiber-coupled photostimulation or a 4-pin male-female Mill-Max connector (ED85100-ND, Mill-Max) for µLED neural probe photostimulation, depending on the type of optical delivery option. The PCB also includes two through-hole pads for battery recharging (at 30 mA for 1 hour). The BLE chip was soldered to the PCB using solder past (4860P-35G, MG Chemicals) and a reflow oven (T-962, puhui), and the other components were hand-soldered using a soldering station (CD-2SQE, JBC).

Following headstage circuit assembly, additional components are needed to operate the headstage with each of the optical delivery modes. For the optical fiber option, a custom fiber ferrule coupler was modeled in FreeCAD (v0.21.0) and 3D-printed in a resin printer (ELEGOO 3 Mars Pro) to fit around the LED on one side and mate with 1.25 mm OD ferrules (Thorlabs) on the other side. The coupler was fixed to the PCB with UV-cure epoxy (Tetric EvoFlow) and covered with liquid black tape (Performix), so that any scattered light was blocked. For the µLED neural probe option, the soldered Mill-Max connector was additionally secured with epoxy (G14250, Thorlabs).

Assembled headstages were weighed with a precision balance (5173, Nahita Blue) and the final dimensions measured with a ruler.

### 4.2. BLE SoC programming

The BLE chip firmware was developed in C using an evaluation board (CYBLE-014008-0-EVAL, Infineon Technologies) and PSoC Creator 4.2 IDE. BLE chips for each headstage were programmed with a programming kit (CYBLE-014008-PROG, Infineon Technologies). The BLE chip was programmed to operate as a slave and once it is turned on, it enters pairing mode. Once paired, it holds in sleep mode until a writing request is received from the BLE transmitter. The BLE chip was programmed to execute the stimulation parameters defined in the app (see section 4.3) and transmitted over the Arduino-compatible transmitter (see section 4.4). The light intensity is controlled using PWM of the 3.5 V supplied by the battery connected to the voltage regulator. If the headstage is used with a µLED neural probe, the maximum voltage is limited in the BLE SoC to 3.2V, using the 3.5V supply as a reference. The chip program also controls the indicative red LED in the headstage according to operational state (on, paired, and unpaired).

### 4.3. App design

A user-friendly standalone application was developed using Visual Studio Community 2022 (v17.6.4) in C#. The app communicates with an Arduino Uno through specific command strings, providing an interface to control and issue AT commands that the Arduino passes to the BLE transmitter.

### 4.4. Transmitter setup

The transmitter setup is composed of a HM-10 BLE Arduino-compatible module and an Arduino Uno rev4. The Arduino code was written in Arduino IDE 2.1.0 to manage Bluetooth communication with the headstage, send writing requests to either assess the battery level or trigger stimulation with the specified parameters, and handling communication with the custom app.

### 4.5. Optical fibers and μLED neural probes fabrication and assessment

The optical fibers were fabricated following an established protocol [44]. The µLED neural probes were fabricated using standard lithography processes in silicon and a custom μLED mounting and passivation process using specially designed 3D printed tools on a pick-and-place machine. The full fabrication process and validation of the neural μLED probes are described in our previous publication [31].

Optical power output of both optical fibers and µLED neural probes coupled to our headstage were taken with a photodiode power sensor (S121C, Thorlabs) connected to a power meter (PM101, Thorlabs). The sensor was placed directly above the light source, at a distance lower than 1 mm. Light sources were powered continuously by the headstage at maximum current.

### 4.6. Battery, operating range, and latency assessment

To assess battery runtime under experimental conditions, the headstage was operated in different modes while the elapsed time until no light was emitted (below 3.5 V or 3.2 V, depending on the optical delivery option) was measured. The operation modes tested were: stand-by (i.e. paired with the emitter, on sleep mode, waiting for trigger); continuous 10 Hz stimulation at maximum intensity, and intermittent 10 Hz (50% duty cycle) protocol with 10 seconds on and 10 second off with an optical fiber and a µLED neural probe.

The assess the maximum distance at which the pairing between the headstage and the transmitter were maintained, pairing was initially performed at a distance of 1 meter and gradually increased by 1 meter steps until connection was lost. The presence of pairing was visually confirmed by observation of the indicator LED on the headstage during this protocol.

To assess system latencies, a camera (FL3-U3-13Y3M-C Flea3 Camera, FLIR) was used to film the full system, from app to headstage, at a frame rate of 150 fps (precision of 6.67 ms). Visual cues were added to the system to indicate when detected trigger events from the PC were first registered in the Arduino, immediately before the transmitter sent a writing request, and finally when the stimulator light was turned on. The difference between the runtime of the code with and without visual cues was calculated to be approximately 30 ms, and this value was subtracted to the measurement of the full system latency.

### 4.7. Mice surgery

Mice (n=2) used for optical fiber tests were Emx1-cre:Ai27D [35], [36], with channelrhodopsin being expressed cortex wide. The mouse used for µLED neural probe tests was injected bilaterally in the ventrolateral periaqueductal gray (vlPAG) (Bregma: AP +4.60, ML +/−0.75, DV −3.0 mm) with a volume of 100 nL of AAV5-CaMKIIa-hChR2(H134R)-EYFP (UNC Gene Therapy) using a Nanoject device (Drummond Scientific; US) in a stereotaxic frame (World Precision Instruments, E06277-002) and under gaseous anaesthesia (sevofluoarane, Sevorane, Abbvie). After injection, the capillary was held in place for 10 minutes to allow volume diffusion and prevent backflow. To allow expression of the virus, behavior was only performed after four weeks.

For optical fiber or µLED implants, mice were anesthetized using sevoflurane (Sevorane, Abbvie) in O2-enriched air, with an induction concentration of 5% and a maintenance concentration of 1.5-2%. Subcutaneous administration of buprenorphine (0.1 mg/kg; Bupaq, Richter Pharma) and 0.9% NaCl (10 mL/kg; 17G15, Advin) were performed prior to the start of the surgery to ensure intra-operative analgesia and hydration, respectively. Mice were placed in a stereotaxic frame (World Precision Instruments, E06277-002) and the dorsal skull was exposed with a midline incision to retract the skin. A burr hole was made in the skull above the M2 (Bregma: AP + 2.50, ML + 1.10) for the optical fiber implantat, or the vlPAG (Bregma: AP +4.60, ML −0.75) for the µLED implant. The skull around the burr hole was covered with a thin layer of OptiBond Solo Plus (29692, Kerr) and cured with UV light. The optic fiber and µLED were then lowered into the burr hole to −0.80 mm and −3.00 mm, to target the M2 and vlPAG, respectively. The burr hole was closed using super glue (64546, UHU) and the implant was secured with OrthoJetBCA dental cement (B1302×6, Lang Dental Mfg). For the µLED implant procedure, operation was checked at this point by turning the µLED on with an external power supply (HY3005D, Kaise). The dental cement surface was painted with white nail polish to avoid light scattering during behavioral testing. Mice were allowed to recover from surgery at least for 3 days.

### 4.8. Behavior testing and analysis

For experiments to confirm wireless headstage control of photostimulation with optical fibers, mice (n=2) were placed in an open-field arena (35 x 35 cm) and allowed to freely explore it for 5 minutes. Then photostimulation was turned on for 1 minute epochs, with an intertrial period of 1 minute where stimulation was off. For each exposure to the arena, 5 stimulation on/off epochs were assessed. Videos were taken with a camera (FL3-U3-13Y3M-C Flea3 Camera, FLIR) placed on top of the arena at 30 fps. Bonsai-RX (v2.7.2) [39] was used to automatically detect and track the animal’s centroid. Total distance traveled and the number of contralateral and ipsilateral turns was calculated with custom python code. Normalization of rotations for each analyzed period was obtained by dividing the number of rotations observed in each period by the total number of rotation in the full session.

For experiments with closed-loop control of photostimulation with the headstage connected to µLED neural probes, one mouse was placed in a custom-made bottomless acrylic open-field arena (35 x 35 cm) on top of a transparent acrylic surface. Videos were recorded from below with a video camera (FL3-U3-13Y3M-C Flea3 Camera, FLIR) at a frame rate of 30 fps. Bonsai-RX software with the DeepLabCut interface [40], [45] running on a computer with an Intel 7 CPU, 64 GB of RAM, and a NVIDIA GeForce RTX 4080 GPU, was used for real-time pose estimation and the automatic classification of translational movement, freezing, and grooming events in video frames from the camera placed at the bottom of the behavioral arena. A DeepLabCut (DLC, v2.2.0) network to estimate the position of 6 animal body parts (paws, snout, and tailbase) was previously trained in a NVIDIA Quadro RTX 6000 GPU with 50 randomly picked frames from 136, 10 minute long, videos of freely moving mice, recorded at 60 fps. The reported train and test errors, for a confidence threshold of 0.90, were 4.04 and 4.22 pixels, respectively, for a frame size of 992 by 896 pixels. This network was loaded in the detect pose node in Bonsai-RX. A python node on Bonsai-RX with custom python code ran a translational movement detector, and freezing and grooming event detectors and counters. Translational movement was detected based on a threshold applied to the cartesian distance travelled by the tailbase between two consecutive frames. Freezing was detected by simultaneously quantifying the stability of the mouse’s body shape (estimated by the polygon area from the 5 body parts) and snout velocity (frame-to-frame distance), for at between 2 frames. Grooming was detected by a decision tree classifier using the following variables from the detected body parts: area between front paws and snout, distance between front paws and snout, and area between back legs and tail base. Previous assessment of the face groom algorithm with data from 5 mice compared to manually scored videos presented a test average accuracy of 97.5%, with an F1 score of 79.3% (data not shown). Detection of translational movement events triggered stimulation requests, via the USB connection to an arduino running the firmata protocol, that sent digital triggers to another arduino running the Bluetooth emitter code that implemented the stimulation parameters/protocol defined in the user app.

The mouse was placed individually into the behavioral arena and allowed to explore freely for a 5 minute period. Following this baseline period, the real-time closed-loop system running in Bonsai started operating, with detection of translational movement triggering µLED activation with blue light pulses (490nm) delivered at a frequency of 10 Hz, 75% intensity, 80% duty cycle, to the vlPAG for a total duration of 10 seconds. To prevent overstimulation, after each stimulation event a 30 seconds refractory period, in which stimulation could not be triggered by translational movement, was implemented. The face groom and freezing classifiers automatically detected these events during behavior and counted the number of events and the time the mice spent in each event.

### 4.9. Statistics

Additional behavioral data analysis and statistics used custom algorithms in Python (v3.8.12) and GraphPad Prism (v10). Statistical tests used can be found in figure legends. Data in graphs is mean ± SEM, except where noted.

## Acknowledgments

The authors would like to thank all members of the Neurophysiology and Neuroengineering lab at FMUP for helpful discussions on device design and implementations. This project was partly funded by “la Caixa” Banking Foundation under grant agreement LCF/PR/HR21-00410 and Portuguese Foundation for Science and Technology (FCT) project 2022.05228.PTDC. MA and MF were supported by FCT doctoral fellowships 2022.14536.BD and 2023.00604.BD, respectively.

## Author Contributions

PS and LJ conceptualized the device. PS designed device circuit, and programmed BLE chip. PS and FN designed the user interface app. PS programmed the communication protocol between app, Arduino, and BLE chip. PS fabricated ferrule coupled optical fibers. MA and TP fabricated µLED neural probes. SR performed viral injections. MF implanted animals with optical fibers and µLED neural probes. PS trained DLC network. PS, SR, and LJ wrote online behavior detection and classification code. PS implemented closed-loop design. MF and SR performed behavior. PS, SR, and LJ analyzed behavior. PM and LJ supervised the work, acquired funding, and provided resources. PS, MF, SR, and LJ wrote the first draft of the manuscript. LJ revised the manuscript. All authors read and approved the final version of the manuscript.

## Ethics statement

All procedures involving animals were conducted in accordance with European Union Directive 2016/63/EU and the Portuguese law on the protection of animals for scientific purposes (DL No 113/2013), and approved by the Portuguese National Authority for Animal Health and the Animal Welfare Committee at FMUP (ORBEA-FMUP).

## Notes

### Competing Interest Statement

The authors have declared no competing interest.

## References

[1] J. G. Bernstein and E. S. Boyden, “Optogenetic tools for analyzing the neural circuits of behavior,” Trends in Cognitive Sciences. 2011, doi: 10.1016/j.tics.2011.10.003.

[2] K. M. Tye and K. Deisseroth, “Optogenetic investigation of neural circuits underlying brain disease in animal models,” Nature Reviews Neuroscience. 2012, doi: 10.1038/nrn3171.

[3] Q. Zhang, T. Li, M. Xu, B. Islam, and J. Wang, “Application of Optogenetics in Neurodegenerative Diseases,” Cellular and Molecular Neurobiology. 2024, doi: 10.1007/s10571-024-01486-1.

[4] K. Deisseroth, “Circuit dynamics of adaptive and maladaptive behaviour,” Nature. 2014, doi: 10.1038/nature12982.

[5] M. R. Warden, J. A. Cardin, and K. Deisseroth, “Optical neural interfaces,” Annual Review of Biomedical Engineering. 2014, doi: 10.1146/annurev-bioeng-071813-104733.

[6] B. A. Humburg et al., “Optogenetic brain-stimulation reward: A new procedure to re-evaluate the rewarding versus aversive effects of cannabinoids in dopamine transporter-Cre mice,” Addict. Biol., 2021, doi: 10.1111/adb.13005.

[7] E. Burguière, P. Monteiro, G. Feng, and A. M. Graybiel, “Optogenetic stimulation of lateral orbitofronto-striatal pathway suppresses compulsive behaviors,” Science (80-.)., vol. 340, no. 6137, pp. 1243–1246, 2013, doi: 10.1126/science.1232380.

[8] Y. Yoon et al., “Neural probe system for behavioral neuropharmacology by bi-directional wireless drug delivery and electrophysiology in socially interacting mice,” Nat. Commun., 2022, doi: 10.1038/s41467-022-33296-8.

[9] Y. Yang et al., “Wireless multilateral devices for optogenetic studies of individual and social behaviors,” Nat. Neurosci., 2021, doi: 10.1038/s41593-021-00849-x.

[10] H. Y. Chen, J. S. Wu, B. Hyland, X. D. Lu, and J. J. J. Chen, “A low noise remotely controllable wireless telemetry system for single-unit recording in rats navigating in a vertical maze,” Med. Biol. Eng. Comput., 2008, doi: 10.1007/s11517-008-0355-6.

[11] P. Silva and L. Jacinto, “Wireless Devices for Optical Brain Stimulation: A Review of Current Developments for Optogenetic Applications in Freely Moving Mice,” Cell. Mol. Bioeng., vol. 18, no. 1, pp. 1–13, Feb. 2025, doi: 10.1007/s12195-024-00832-z.

[12] B. A. Wester, R. H. Lee, and M. C. LaPlaca, “Development and characterization of in vivo flexible electrodes compatible with large tissue displacements.,” J. Neural Eng., vol. 6, no. 2, p. 024002, 2009, doi: 10.1088/1741-2560/6/2/024002.

[13] M. A. Rossi, V. Go, T. Murphy, Q. Fu, J. Morizio, and H. H. Yin, “A wirelessly controlled implantable LED system for deep brain optogenetic stimulation,” Front. Integr. Neurosci., 2015, doi: 10.3389/fnint.2015.00008.

[14] K. L. Montgomery et al., “Wirelessly powered, fully internal optogenetics for brain, spinal and peripheral circuits in mice,” Nat. Methods, 2015, doi: 10.1038/nmeth.3536.

[15] G. Shin et al., “Flexible Near-Field Wireless Optoelectronics as Subdermal Implants for Broad Applications in Optogenetics,” Neuron, 2017, doi: 10.1016/j.neuron.2016.12.031.

[16] Y. Zhao et al., “Wirelessly Operated, Implantable Optoelectronic Probes for Optogenetics in Freely Moving Animals,” IEEE Trans. Electron Devices, 2019, doi: 10.1109/TED.2018.2882397.

[17] L. Li et al., “Colocalized, bidirectional optogenetic modulations in freely behaving mice with a wireless dual-color optoelectronic probe,” Nat. Commun., 2022, doi: 10.1038/s41467-022-28539-7.

[18] S. Il Park et al., “Ultraminiaturized photovoltaic and radio frequency powered optoelectronic systems for wireless optogenetics,” J. Neural Eng., 2015, doi: 10.1088/1741-2560/12/5/056002.

[19] R. Dagnew et al., “CerebraLux: a low-cost, open-source, wireless probe for optogenetic stimulation,” Neurophotonics, 2017, doi: 10.1117/1.nph.4.4.045001.

[20] S. Anpilov et al., “Wireless Optogenetic Stimulation of Oxytocin Neurons in a Semi-natural Setup Dynamically Elevates Both Pro-social and Agonistic Behaviors,” Neuron, 2020, doi: 10.1016/j.neuron.2020.05.028.

[21] S. Orguc, J. Sands, A. Sahasrabudhe, P. Anikeeva, and A. P. Chandrakasan, “Modular Optoelectronic System for Wireless, Programmable Neuromodulation during Free Behavior,” 2020, doi: 10.1109/EMBC44109.2020.9175600.

[22] J. Park et al., “A wireless, solar-powered, optoelectronic system for spatial restriction-free long-term optogenetic neuromodulations,” Sci. Adv., 2023, doi: 10.1126/sciadv.adi8918.

[23] L. M. Fischer et al., “Gold cleaning methods for electrochemical detection applications,” Microelectron. Eng., vol. 86, no. 4–6, pp. 1282–1285, Apr. 2009, doi: 10.1016/j.mee.2008.11.045.

[24] C. Y. Kim et al., “Soft subdermal implant capable of wireless battery charging and programmable controls for applications in optogenetics,” Nat. Commun., 2021, doi: 10.1038/s41467-020-20803-y.

[25] W. Chen et al., “The Roles of Optogenetics and Technology in Neurobiology: A Review,” Frontiers in Aging Neuroscience. 2022, doi: 10.3389/fnagi.2022.867863.

[26] S. Il Park et al., “Stretchable multichannel antennas in soft wireless optoelectronic implants for optogenetics,” Proc. Natl. Acad. Sci. U. S. A., 2016, doi: 10.1073/pnas.1611769113.

[27] P. Gutruf et al., “Fully implantable optoelectronic systems for battery-free, multimodal operation in neuroscience research,” *Nat*. Electron., 2018, doi: 10.1038/s41928-018-0175-0.

[28] S. T. Lee, P. A. Williams, C. E. Braine, D. T. Lin, S. W. M. John, and P. P. Irazoqui, “A Miniature, Fiber-Coupled, Wireless, Deep-Brain Optogenetic Stimulator,” IEEE Trans. Neural Syst. Rehabil. Eng., 2015, doi: 10.1109/TNSRE.2015.2391282.

[29] A. Nourizonoz et al., “EthoLoop: automated closed-loop neuroethology in naturalistic environments,” Nat. Methods, 2020, doi: 10.1038/s41592-020-0961-2.

[30] J. F. Schweihoff, M. Loshakov, I. Pavlova, L. Kück, L. A. Ewell, and M. K. Schwarz, “DeepLabStream enables closed-loop behavioral experiments using deep learning-based markerless, real-time posture detection,” *Commun*. Biol., 2021, doi: 10.1038/s42003-021-01654-9.

[31] M. Abrantes et al., “Small form factor implantable neural probe with efficient flip chip µLED for in vivo optogenetics,” Biomed. Microdevices, 2025, doi: 10.1007/s10544-025-00754-1.

[32] V. Emiliani et al., “Optogenetics for light control of biological systems,” Nat. Rev. Methods Prim., 2022, doi: 10.1038/s43586-022-00136-4.

[33] S. F. Owen, M. H. Liu, and A. C. Kreitzer, “Thermal constraints on in vivo optogenetic manipulations,” Nat. Neurosci., 2019, doi: 10.1038/s41593-019-0422-3.

[34] N. Grossman, K. Nikolic, C. Toumazou, and P. Degenaar, “Modeling study of the light stimulation of a neuron cell with channelrhodopsin-2 mutants,” IEEE Trans. Biomed. Eng., 2011, doi: 10.1109/TBME.2011.2114883.

[35] J. A. Gorski, T. Talley, M. Qiu, L. Puelles, J. L. R. Rubenstein, and K. R. Jones, “Cortical excitatory neurons and glia, but not GABAergic neurons, are produced in the Emx1-expressing lineage,” J. Neurosci., vol. 22, no. 15, pp. 6309–6314, 2002, doi: 10.1523/jneurosci.22-15-06309.2002.

[36] L. Madisen et al., “A toolbox of Cre-dependent optogenetic transgenic mice for light-induced activation and silencing,” Nat. Neurosci., vol. 15, no. 5, pp. 793–802, 2012, doi: 10.1038/nn.3078.

[37] L. A. V. Magno et al., “Optogenetic stimulation of the M2 cortex reverts motor dysfunction in a mouse model of Parkinson’s disease,” J. Neurosci., 2019, doi: 10.1523/JNEUROSCI.2277-18.2019.

[38] V. Gradinaru, M. Mogri, K. R. Thompson, J. M. Henderson, and K. Deisseroth, “Optical deconstruction of parkinsonian neural circuitry,” Science (80-.)., 2009, doi: 10.1126/science.1167093.

[39] G. Lopes et al., “Bonsai: An event-based framework for processing and controlling data streams,” Front. Neuroinform., vol. 9, no. APR, 2015, doi: 10.3389/fninf.2015.00007.

[40] G. A. Kane, G. Lopes, J. L. Saunders, A. Mathis, and M. W. Mathis, “Real-time, low-latency closed-loop feedback using markerless posture tracking,” Elife, 2020, doi: 10.7554/ELIFE.61909.

[41] P. Tovote et al., “Midbrain circuits for defensive behaviour,” Nature, 2016, doi: 10.1038/nature17996.

[42] C. E. Vaaga, S. T. Brown, and I. M. Raman, “Cerebellar modulation of synaptic input to freezing-related neurons in the periaqueductal gray,” Elife, 2020, doi: 10.7554/eLife.54302.

[43] J. Signoret-Genest, N. Schukraft, S. L. Reis, D. Segebarth, K. Deisseroth, and P. Tovote, “Integrated cardio-behavioral responses to threat define defensive states,” Nat. Neurosci., 2023, doi: 10.1038/s41593-022-01252-w.

[44] D. R. Sparta, A. M. Stamatakis, J. L. Phillips, N. Hovelsø, R. Van Zessen, and G. D. Stuber, “Construction of implantable optical fibers for long-term optogenetic manipulation of neural circuits,” Nat. Protoc., 2012, doi: 10.1038/nprot.2011.413.

[45] A. Mathis et al., “DeepLabCut: markerless pose estimation of user-defined body parts with deep learning,” Nat. Neurosci., vol. 21, no. 9, pp. 1281–1289, 2018, doi: 10.1038/s41593-018-0209-y.

